# A Pathway and Network Oriented Approach to Enlighten Molecular Mechanisms of Type 2 Diabetes Using Multiple Association Studies

**DOI:** 10.1101/2020.01.14.905547

**Authors:** Burcu Bakir-Gungor, Miray Unlu Yazici, Gokhan Goy, Mustafa Temiz

## Abstract

Diabetes Mellitus (DM) is a group of metabolic disorder that is characterized by pancreatic dysfunction in insulin producing beta cells, glucagon secreting alpha cells, and insulin resistance or insulin in-functionality related hyperglycemia. Type 2 Diabetes Mellitus (T2D), which constitutes 90% of the diabetes cases, is a complex multifactorial disease. In the last decade, genome-wide association studies (GWASs) for type 2 diabetes (T2D) successfully pinpointed the genetic variants (typically single nucleotide polymorphisms, SNPs) that associate with disease risk. However, traditional GWASs focus on the ‘the tip of the iceberg’ SNPs, and the SNPs with mild effects are discarded. In order to diminish the burden of multiple testing in GWAS, researchers attempted to evaluate the collective effects of interesting variants. In this regard, pathway-based analyses of GWAS became popular to discover novel multi-genic functional associations. Still, to reveal the unaccounted 85 to 90% of T2D variation, which lies hidden in GWAS datasets, new post-GWAS strategies need to be developed. In this respect, here we reanalyze three meta-analysis data of GWAS in T2D, using the methodology that we have developed to identify disease-associated pathways by combining nominally significant evidence of genetic association with the known biochemical pathways, protein-protein interaction (PPI) networks, and the functional information of selected SNPs. In this research effort, to enlighten the molecular mechanisms underlying T2D development and progress, we integrated different in-silico approaches that proceed in top-down manner and bottom-up manner, and hence presented a comprehensive analysis at protein subnetwork, pathway, and pathway subnetwork levels. Our network and pathway-oriented approach is based on both the significance level of an affected pathway and its topological relationship with its neighbor pathways. Using the mutual information based on the shared genes, the identified protein subnetworks and the affected pathways of each dataset were compared. While, most of the identified pathways recapitulate the pathophysiology of T2D, our results show that incorporating SNP functional properties, protein-protein interaction networks into GWAS can dissect leading molecular pathways, which cannot be picked up using traditional analyses. We hope to bridge the knowledge gap from sequence to consequence.

## 1 Introduction

More than 400 million adults struggle with Diabetes Mellitus, and this number is expected to reach 600 million by 2040 (International Diabetes Federation, 2017). Type 1 and Type 2 Diabetes Mellitus (T1DM, T2DM) are the two main types of Diabetes, which contribute to worldwide health care problem by not properly using blood glucose for energy in the body. While T1DM is mostly related with pancreatic beta cell damage, T2DM is both associated with beta cells’ functionality and insulin resistance (DeFronzo et al., 2015; Zheng et al., 2018). Recently, with the help of antidiabetic agents, significant progress has been made in maintaining the glycemic control (blood sugar level) in T2D patients. Still, the targeted glycated hemoglobin levels could not be maintained for 40% of the adults with diabetes in USA. The decrease in pancreatic beta cell functionality and the increase in the insulin sensitivity of T2D patients over the time, eventually gave rise to the imbalance of A1C level and antidiabetic treatment gap (Freeman, 2013). This kind of imbalance and dysfunctionality emerges as a result of the complex interactions among the environmental and genetic risk factors. In this respect, the etiology, driving factors and the genetic predispositions responsible for the increased susceptibility of T2D needed to be well understood in developing new drugs and treatments for this disorder. In this kind of complex diseases, the investigations of different mechanisms of actions may provide benefits for therapeutic approaches. Therefore, post analysis of high throughput studies conducted at different molecular levels and the elucidation of targeted genes and pathways associated with T2D are crucial.

The widespread introduction of large-scale genetic studies has enabled researches to investigate the genetic frameworks of complex disorders. During the last decade, genome wide association studies (GWAS) are widely used to identify the risk factors of complex diseases, to better understand the biological mechanisms of these diseases, and hence to help the discovery of novel therapeutic targets (Claussnitzer et al., 2020). Despite GWASs has led to a remarkable range of discoveries in human genetics (Visscher et al., 2017), it has some shortcomings. One important shortcoming of GWAS stems from its testing each marker once at a time for association with disease. Since these studies evaluate the significance of the variants individually, they probably miss the SNPs that have low contribution to disease individually, but might be important when interacting collectively. Moreover, in traditional GWASs, the functional effects of significant SNPs, predicted at the splicing, transcriptional, translational, and post-translational levels are usually neglected. Although GWAS identified more than 140 independent loci influencing the risk of T2D (Bonàs-Guarch et al., 2018; Mahajan et al., 2018b, 2018a; Mercader and Florez, 2017; Scott et al., 2017; Xue et al., 2018; Zhao et al., 2017), most of these loci are driven by common variants and the mechanistic understanding has only been achieved only for a couple of these genes. In this respect, post-GWAS strategies need to be developed to enlighten the molecular mechanisms underlying T2D development and progress (White et al., 2019).

Recent studies indicated that the methods focusing on pathways rather than individual genes can detect significant coordinated changes since these genes act in a synergistic mode in a biological pathway (Nguyen et al., 2019). Pathway analysis can hypothetically improve power to uncover genetic factors relevant to disease mechanisms, because identifying the accumulation of small genetic effects acting in a common pathway is often easier than mapping the individual genes within the pathway that contribute to disease susceptibility remarkably (Kao et al., 2017; Lamparter et al., 2016; Thrash et al., 2019). The profound discovery that T2D is genetically heterogeneous suggested that the genetic defects might converge on common pathways building up the final similar phenotype. Besides providing the opportunity to investigate additional therapies that reverse the effects of a particular genetic defect, these findings also may encourage scientist to understand the aberrant networks at genetic, cellular and physiological levels and to devise pharmacological and nonpharmacological intervention strategies.

Inspired by these findings, in this study, we reanalyzed three meta GWAS dataset of T2D, using the methodology that we have developed to identify disease-associated pathways by combining nominally significant evidence of genetic association with the known biochemical pathways, protein-protein interaction (PPI) networks, and the functional information of selected SNPs (Bakir-Gungor et al., 2014).

## 2 Materials and Methods

### 2.1 Datasets

#### 2.1.1 70K for T2D Meta-analysis data (T2D1)

Bonàs-Guarch *et. al.* collected T2D genome wide association study (GWAS) data, representing 12,931 cases and 57,196 controls of European ancestry from EGA and dbGaP databases (Bonàs-Guarch et al., 2018). In 70KforT2D meta-analysis data, each dataset was quality controlled and each cohort was imputed to reference panels (1000G and UK10K). Variants which were selected for IMPUTE2 info score ≥ 0.7, MAF ≥ 0.001 and, Hardy-Weinberg equilibrium (HWE) controls p > 1×10-6, were meta-analyzed. For more details about the followed quality control procedure and association analysis of 70KforT2D dataset, please see (Bonàs-Guarch et al., 2018).

#### 2.1.2 Meta-analysis of DIAGRAM, GERA, UKB GWAS datasets (T2D2)

Xue *et. al.* performed a meta-analysis of GWAS in T2D by gathering DIAGRAM, GERA, UKB GWAS datasets (Xue et al., 2018). 62,892 cases and 596,424 controls of European ancestry in total were obtained after quality controls and imputed to 1000 Genomes Project. Linkage disequilibrium (LD) score regression analysis was demonstrated. Variants were filtered for GERA and UKB using IMPUTE2 info score ≥ 0.3, MAF ≥ 0.01, HWE controls p > 1×10-6. Further details about DIAGRAM imputed data in stages 1 and 2, genotyping, quality control and association analysis for each dataset can be found in (Xue et al., 2018).

#### 2.1.3 Type 2 Diabetes GWAS Meta-analysis Dataset #3 (T2D3)

Mahajan *et. al.* collected T2D GWAS datasets from 32 studies including 74,124 cases and 824,006 controls of European population, and aggregated data after initial analyses (Mahajan et al., 2018a). Following quality control checks, the imputation of studies was performed using Haplotype Reference Consortium reference panel, except for deCODE GWAS, where population-specific reference panel was used for imputation. For detailed information, please refer to (Mahajan et al., 2018a).

#### 2.1.4 Protein-protein interaction (PPI) dataset

A human protein-protein interaction (PPI) network (interactome data) containing 13,460 proteins and 141,296 protein-protein interactions was derived from (Ghiassian et al., 2015) and used in subnetwork identification steps of this study.

### 2.2 Methods

To enlighten the molecular mechanisms underlying T2D development and progress, here we integrated different in-silico approaches that proceed in top-down manner and bottom-up manner, as summarized in Figure 1. Via combining nominally significant evidence of genetic association with the known biochemical pathways, PPI networks, and the functional information of selected SNPs, our proposed approach identifies disease-associated pathways.

**Figure 1.**
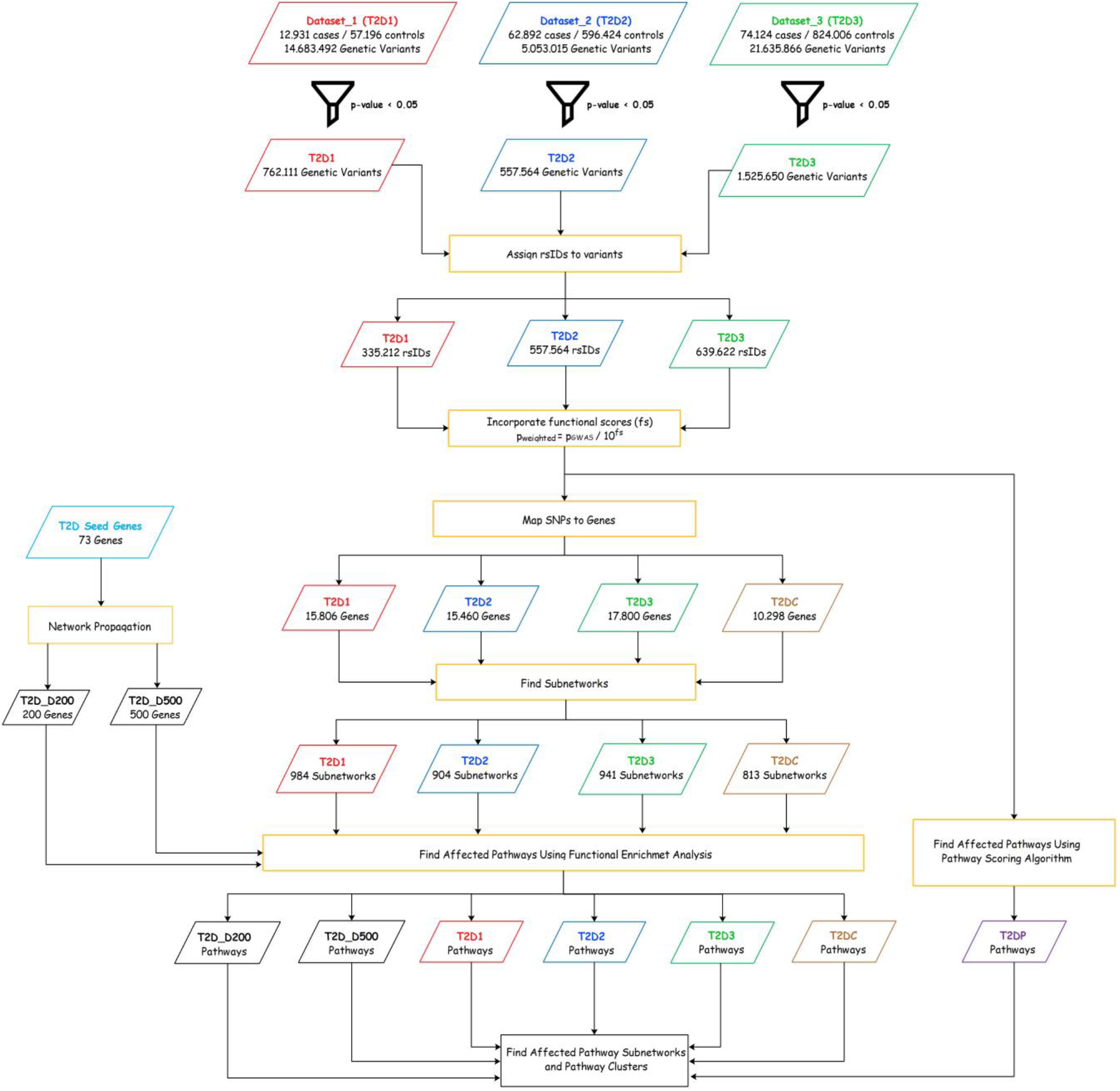
Summary of our pathway and network oriented approach to enlighten T2D mechanisms using multiple association studies.

#### 2.2.1 Preprocessing

Association summary statistics for the T2D1, T2D2, T2D3 datasets were downloaded from each project’s website. This summary statistics data includes i) marker name as chromosome and position, ii) effect allele, iii) non-effect allele, and iv) p-value of association. To be able to assess the collective effect of the variants detected in GWAS with mild effects, all variants were filtered using p<0.05 cutoff, as suggested in previous studies (Bakir-Gungor et al., 2013, 2015; Bakir-Gungor and Sezerman, 2011, 2013; Baranzini et al., 2009).

#### 2.2.2 Assigning rsIDs to identified SNPs

While T2D2 dataset provides associated rsIDs of the identified SNPs in the summary statistics data, T2D1 and T2D3 datasets only provide chromosome and position information as marker name of the variants and do not provide associated rsIDs. In this respect, fast and easy variant annotation protocol introduced by (Yang and Wang, 2015) is utilized to assign associated rsIDs to the identified SNPs using hg19 or hg38 reference genomes, depending on the provided genomic coordinates at T2D1, T2D3 datasets.

#### 2.2.3 Assessing the Functional Impacts of Genetic Variants

To assess the functional impact of a non-synonymous change on proteins, numerous computational methods have been developed, as reviewed in (Zeng and Bromberg, 2019). These methods can be classified as following: i) methods that score mutations on the basis of biological principles, ii) methods that use existing knowledge about the functional effects of mutations in the form a training set for supervised machine learning (Carter et al., 2013). Most of these methods assign a numeric score to the non-synonymous change, indicating the predicted functional impact of an amino acid substitution. To identify likely functional missense mutations, Douville *et. al.* developed a tool called The Variant Effect Scoring Tool (VEST), that utilizes Random Forest as a supervised machine learning algorithm (Douville et al., 2016). Douville *et. al.* represents all mutations with a set of 86 quantitative features; and used missense variants from the Human Gene Mutation Database as a positive class and common missense variants detected in the Exome Sequencing Project (ESP) as a negative class, in their training set (Douville et al., 2016). Since VEST scores result in 0.9 sensitivity and 0.9 specificity values, these scores are utilized to assess the functional impacts of genetic variants in our study.

#### 2.2.4 Assigning SNPs to genes

Several post-GWAS studies map disease-associated SNPs to genes based on physical distance (Segrè et al., 2010), linkage disequilibrium (LD) (Pers et al., 2015), or a combination of both (Wood et al., 2014). In this respect, to aggregate SNP summary statistics into gene scores, several methods have been proposed (Li et al., 2011; Liu et al., 2010; Segrè et al., 2010). Via applying inverse chi-squared quantile transformation on SNP p-values, most of these methods firstly calculate chi-squared values. Secondly, within a window encompassing the gene of interest, some of these methods focus only on the most significant SNP, and assign the maximum-of-chi-squares as the gene score statistic (Lee et al., 2011; Segrè et al., 2010). Some other methods combine results for all SNPs in the gene region by using the sum-of-chi-squares statistic (Wang et al., 2011). In order to compute a well-calibrated p-value for the statistic, gene size and LD structure correction is also critical. (Lamparter et al., 2016) rigorously analyzed the effects of using the sum and the maximum of chi-squared statistics, which correspond to the strongest and the average association signals per gene, respectively. (Lamparter et al., 2016) proposed a fast and efficient methodology, Pascal, that calculates gene scores by aggregating SNP p-values from a GWAS meta-analysis (without the need for individual genotypes), while correcting for LD structure. Pascal only requires SNP-phenotype association summary statistics and do not require genotype data. Hence, we utilized this tool in our study to map SNPs into genes.

#### 2.2.5 The Identification of Dysregulated Modules

High throughput experiments enable us to gain better understanding of the functions of the biological molecules in the cell. In addition to the individual activities of these molecules, the molecular interactions are essential to elucidate these molecular mechanisms. In this regard, human protein-protein interaction (PPI) networks represent the interactions between human proteins. Via analyzing PPI networks, specific sets of proteins (modules) associated with disease phenotype could be detected. This idea is exploited in several post-GWAS analyzes (Bakir-Gungor et al., 2013, 2014, 2015; Bakir-gungor and Sezerman, 2013; Bakir-Gungor and Sezerman, 2011; Chang et al., 2018).

An undirected graph could be defined as G = (V, E), in which the vertex or nodes (V) represent proteins, edges (E) represent the physical interactions among proteins, and graph (G) represent protein-protein interaction (PPI) network. A group of proteins in a PPI network that works together to carry out a specific set of functions can be defined as a subnetwork. With the idea of proteins working as a team, disease related protein subnetwork detection has been widely investigated. Active subnetwork search algorithms are originally proposed to identify dysregulated modules in a PPI via utilizing the gene expression values measured in a microarray study (Ideker et al., 2002). p-values of the genes indicate the significance of expression changes of a gene over certain conditions are mapped to PPI and a search algorithm identifies dysregulated modules. Our group and several others later extended this idea to post-GWAS analyzes, where the SNPs are initially mapped to genes and then the p-values of a gene (genotypic p-values) indicate the significance of a gene in the genetic association study. In this study, to detect dysregulated modules, we use the following two approaches that proceed in top-down and bottom-up manners.

##### 2.2.5.1 Using Subnetwork Identification Algorithms (Top-down approach)

The methodology proposed by (Ideker et al., 2002) to identify active modules in PPI networks, became a pioneer study in this field. While this method brings together the nodes that are highly affected by the condition under study, it also gives a chance to the neighbor nodes of these highly affected nodes, even if they are not highly affected. In this method, firstly a scoring function is defined for each subnetwork and then the problem turned into a search problem of a subnetwork, which maximizes this score. More specifically, to score a subnetwork, the genotypic p-value is converted to a z-score using the equation below, where Φ^ (−1) indicates inverse normal probability distribution.

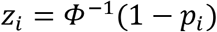

The total z score (z_A_) of the subnetwork A, including k genes is calculated as follows:

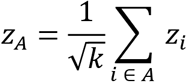

While this score is normalized using the following equation, where μ and σ indicates mean and standard deviation, respectively; the subnetwork scores are also calibrated by the Monte Carlo method.

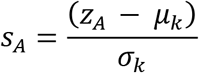

Once the subnetwork score is defined, greedy approach, genetic algorithm, and simulated annealing are popular search strategies in active subnetwork identification methodologies. In this study, greedy approach is used during the search steps of the algorithm, and the subnetwork score cutoff is chosen as 3, as suggested in the original paper (Ideker et al., 2002) to select biologically meaningful subnetworks.

##### 2.2.5.2 Using Network Propagation (Bottom Up Approach)

Based on the idea that the disease-related proteins do not concentrate in a specific region, studies focus on the estimation of dysregulated modules by using the degree of affected nodes information and edges (protein interaction). (Ghiassian et al., 2015) proposed DIseAse MOdule Detection (DIAMOnD) algorithm that finds out dysregulated modules by adding other possible proteins around the known disease protein clusters. Based on random walking, a defined walker starts from a random seed protein and moves through other nodes along the connections of the network. It is hypothesized that more frequently visited proteins are closer to seed proteins (proteins that are known to be associated with the disease). The probability of a random protein with k interaction having k_s_ interaction with seed proteins is calculated by the hyper-geometric distribution as follows:

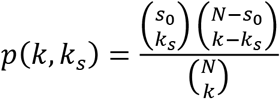

Here, N denotes the number of proteins, s_0_ denotes the number of seed proteins associated with a particular disease. Whether a protein in the PPI network is randomly interact with the seed protein is calculated by the p-value in equation below. In this way, initiating from seed proteins, other candidate proteins associated with the disease can be identified.

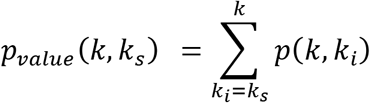

#### 2.2.6 Functional Enrichment

In multifactorial complex disorders, a single factor is unlikely to explain the disease mechanism. Within this scope, functional enrichment analysis focuses on interconnection of terms and functional groups in networks to predict affected pathways for the interested disease. Hyper geometric test and correction methods such as Bonferroni and Benjamini-Hoschberg are used for analyses. Hyper geometric p-value determines the significance of gene enrichment above a certain threshold form predefined functional terms.

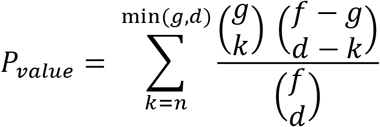

Accordingly, important pathways in the disease and upregulated and downregulated target genes in the pathway are predicted and given as output. In this study, ClueGO (Bindea et al., 2009) is utilized for enrichment analysis. KEGG biological pathways are used as reference pathways.

#### 2.2.7 Construction of Pathway Network

Figure 2 summarizes our steps regarding pathway-pathway biological network generation and pathway subnetwork identification. In order to establish a pathway network, first of all, the relationships between the genes and 288 KEGG biological pathways need to be analyzed. This relationship is revealed via examining whether the gene of interest is found in a specific pathway or not. For example, if pathway i includes gene j, a value of 1 is assigned to index_i,j_ in the gene-term matrix and if not, a value of 0 is given to this index. Hence, the created gene-term matrix is a binary matrix, as shown in Figure 2. Secondly, the relationships between pathways need to be analyzed. For this purpose, the term - term matrix is formed by using the previously obtained gene - term matrix, as illustrated in Figure 2. Kappa score metric is used to determine the relationships among the pathways. The equation expressing the Kappa score for any two pathways A, B is given as follows:

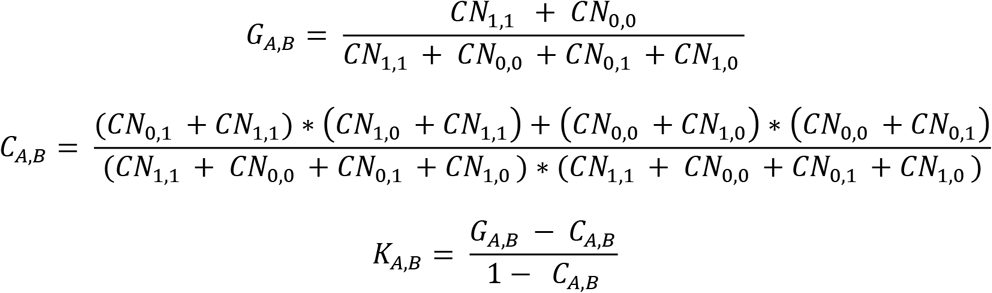

where, G_A,B_ represents the observed contingency, C_A,B_ represents random contingency and K_A,B_ represents the Kappa score between pathways A and B. CN_1,1_, CN_0,0_, CN_1,0_, CN_0,1_ counters are calculated as following. If the gene of interest is present in both compared pathways, CN_1,1_ counter is increased by 1. Following the same idea, the values of other counters are calculated. Kappa scores, which express the relationships between pairs of pathways, was obtained using observed contingency (G) and random contingency (C) values and stored in term - term matrix. Via applying a threshold on Kappa scores, human KEGG pathway network is created. The pathway network generation steps are implemented in Java.

**Figure 2.**
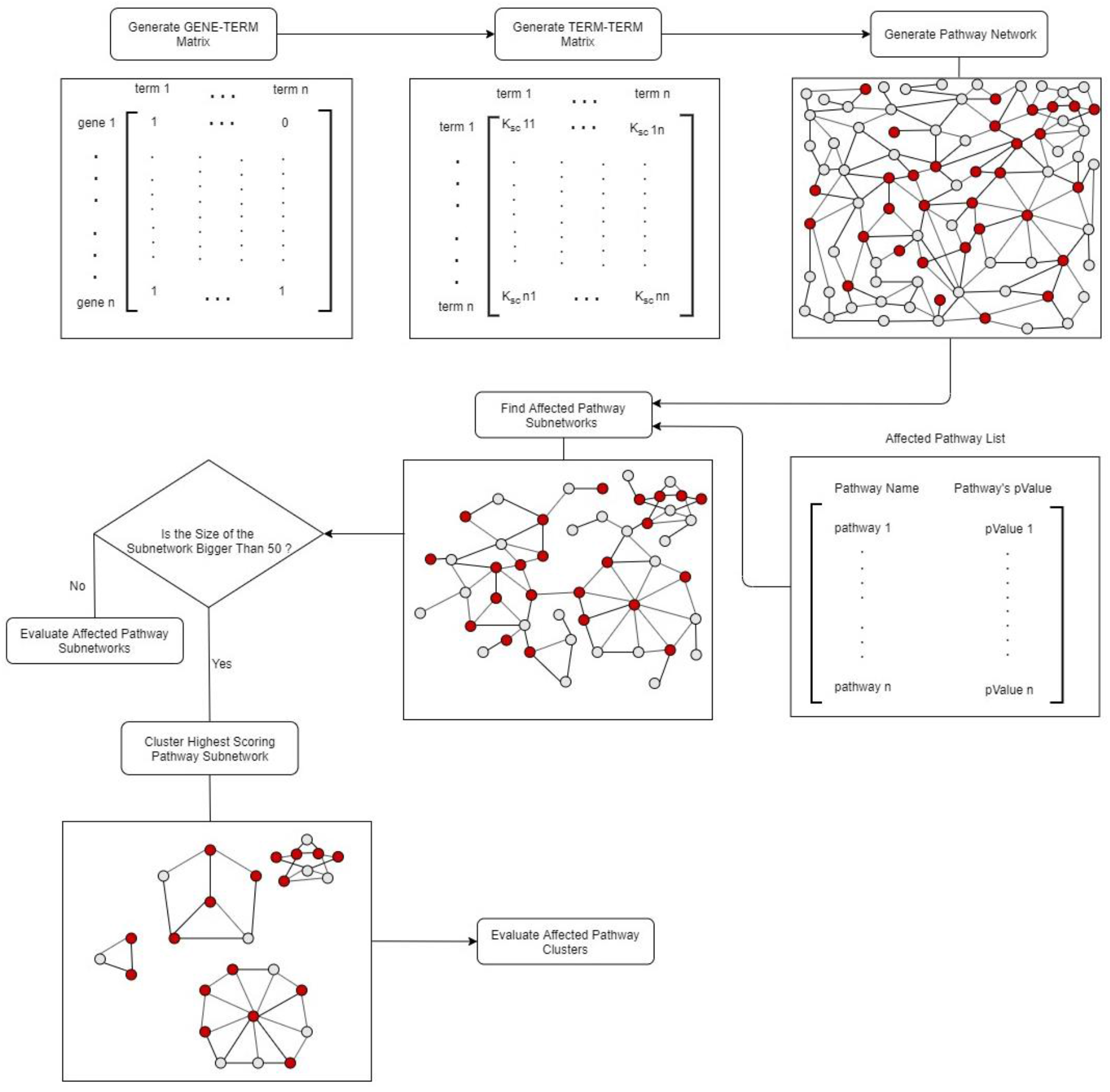
Flowchart of pathway network generation and pathway subnetwork identification.

#### 2.2.8 The Identification of Affected Pathway Subnetworks and Pathway Clusters

To be able to utilize the interrelated structure of the pathways, we proposed to apply subnetwork identification methodologies on the generated pathway networks, hence disease related affected pathway subnetworks could be identified. A classical subnetwork identification algorithm requires the following two information: i) the biological network file, ii) significance of the nodes. In the regular subnetwork identification problem, while (i) refers to a PPI network, (ii) refers to the significance values of the genes, obtained in a microarray experiment. Here, for (i), we used the pathway network that we generated as described in Section 2.2.7. Regarding (ii), the functional enrichment step, as explained in Section 2.2.6 outputs affected pathway lists with their p-values, indicating the importance of a pathway for the phenotype under study. Hence, to obtain the affected pathway subnetworks, a similar methodology, as described in Section 2.2.5.1 is followed. Instead of using a protein-protein interaction network, in this step, the generated pathway network, as explained in Section 2.2.7, is used. Instead of using the significance values of the proteins, in this step, the significance values of the pathways, generated in Functional Enrichment Step, Section 2.2.6, is used. To select biologically meaningful subnetworks among all generated subnetworks, the subnetwork score cutoff is chosen as 3, as suggested in the original paper (Ideker et al., 2002). If the size of the identified subnetwork is bigger than 50, this pathway subnetwork is further sub-divided to find disease related pathway clusters. At this step, we used a graph theoretic clustering algorithm, Molecular Complex Detection (MCODE) to discover densely connected pathway clusters in the T2D affected pathway subnetwork (Bader and Hogue, 2003). In order to confine the dense regions in a PPI, MCODE exploits vertex weighting by local neighborhood density and outward traversal from a locally dense seed protein. In our problem setting, while the PPI refers to the generated pathway network, proteins refer to the pathways. The advantage of MCODE over other graph clustering methods is its allowance for the i) fine-tuning of clusters of interest without considering the rest of the network and ii) inspection of cluster interconnectivity, which is relevant for pathway networks (Bader and Hogue, 2003). It uses 4 different parameters to find clusters: cut off value, K-core value, haircut and fluff parameters. The cut off value sets the intensity of the cluster to be estimated. The K-core parameter allows to assign weights to the nodes, which is later used by MCODE to reduce the running time complexity. The haircut parameter, which is a binary parameter, allows the elimination of nodes considered to be topologically irrelevant. The fluff parameter allows someone to set the size of the cluster, which is estimated topologically in the default mode (Bader and Hogue, 2003). In our analyses, the default values of these parameters are used. In the last step, the identified T2D affected pathway subnetworks and pathway clusters are evaluated.

#### 2.2.9 Pathway Scoring Algorithm

Integration of SNPs across genes and pathways in GWASs has potential to make significant advancement in statistical power and in enlightening relevant biological mechanisms. However, this process is challenging because of the multi-functional roles of genes in several biological processes and the inadequate information about all phenotype – process pairs. In this regard, Pascal (Pathway scoring algorithm) is a robust tool to calculate gene and pathway scores from SNP-phenotype association summary statistics (Lamparter et al., 2016). It does not require genotype data. Firstly, they calculate gene scores by aggregating SNP p-values from a GWAS meta-analysis, and also by correcting for LD structure. While computing the gene scores, they compared the effect of using the sum of chi-squared statistics (average association signals per gene) with the effect of using max of chi-squared statistics (strongest association signals per gene) (Lamparter et al., 2016). Secondly, they calculate pathway scores via aggregating the scores of genes that belong to the same pathways by using modified Fisher method (Lamparter et al., 2016).

#### 2.2.10 Comparison of the Identified Subnetworks and Pathways from Different Datasets Using Normalized Mutual Information (NMI)

In order to evaluate the similarities between two different community detection algorithms, (Xuan Vinh et al., 2010) and (Tripathi et al., 2016) proposed to use Normalized Mutual Information. Let U and V be the sets of subnetworks that are identified using different datasets. Let U= {U_1_, …., U_R_} denote the set of R different subnetworks identified using dataset x, and let V= {V_1_, …., V_S_} denote the set of S different subnetworks identified using dataset y. The following contingency table (Table 1) illustrates the numbers of shared genes between pairs of subnetworks. In other words, n_ij_ indicates the number of common genes between subnetworks U_i_ and V_j_. The entropy of communities H(U), H(V) and mutual information I (U, V) are calculated as following.

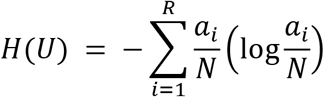

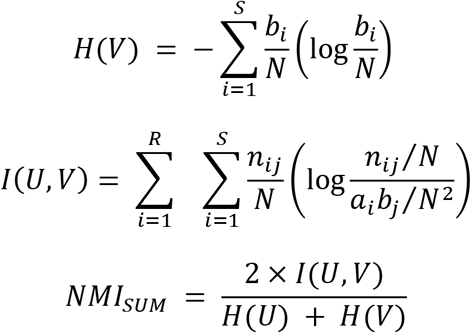

**Table 1.** Contingency table of overlapping genes (n_i, j_) between subnetworks U_i_ and V_j_, where U and V indicate the sets of subnetworks identified via using datasets X and Y, respectively.

| U \| V | V<sub>1</sub> | V<sub>2</sub> | ... | V<sub>S</sub> | Sum |
| --- | --- | --- | --- | --- | --- |
| U<sub>1</sub> | n<sub>11</sub> | n<sub>12</sub> | ... | n<sub>1S</sub> | a<sub>1</sub> |
| U<sub>2</sub> | n<sub>21</sub> | n<sub>22</sub> | ... | n<sub>2S</sub> | a<sub>2</sub> |
| ... | ... | ... | ... | ... | ... |
| U<sub>R</sub> | n<sub>R1</sub> | n<sub>R2</sub> | ... | n<sub>RS</sub> | a<sub>R</sub> |
| Sum | b<sub>1</sub> | b<sub>2</sub> | ... | b<sub>S</sub> | N |

Here, I (U, V) indicates the amount of information shared between U and V communities. NMI_SUM_ is used to compare the clusters in the range of [0,1], where the value 0 refers no similarity between clusters (Vinh et al., 2010).

## 3 Results

Based on the idea that the genes and proteins perform cellular functions in a coordinated fashion, understanding the co-operations of proteins in interaction networks may help to identify candidate biomarkers. In this study, we proposed an integrative approach that concurrently analyzes multiple association studies, the functional impacts of these variants, incorporates the interaction partners of susceptibility genes, detects a pathway network of functionally enriched pathways and finally determines the clusterings and subnetworks of affected pathways. The methodology proposed in Figure 1 is applied on three meta-analyses of GWAS data, which are introduced in Section 2.1. As summarized in Table 2, T2D1, T2D2 and T2D3 datasets include 14 .683.492, 5.053.015, 21.635.866 SNPs respectively. After the filtration of 3 GWAS datasets using p< 0.05 cutoff, the SNPs with mild effects are collected and the numbers of genetic variants are reduced to 762,111, 557,564 and 1,525,650, for T2D1, T2D2 and T2D3 datasets, respectively. Chromosomal position, reference allele, altered allele information of genetic variants are utilized to assign rsIDs. 335,212 and 639,622 rsIDs are assigned to T2D1 and T2D3 datasets, as explained in Section 2.2.2 (Reference genome: hg19). 557,564 rsIDs presented as part of T2D2 dataset is used for further analyses. In the next step, functional scores are assigned to each SNP via using VEST (Douville et al., 2016), as explained in Section 2.2.3. Weighted p-values (pW) are calculated for SNPs via combining the genetic association p-values with functional scores (FS) p_w_=pGWAS/10^FS^, as proposed by (Saccone et al., 2008). Then, SNPs are mapped to 15,806, 15,460 and 17,200 genes for T2D1, T2D2 and T2D3 datasets, respectively. Combined p-values of 10,298 common genes among three datasets are calculated using Fisher’s combined test (Fisher, 1934), and called as T2D-combined (T2DC) in the rest of this paper. For the detection of dysregulated modules, top-down and bottom-up approaches are followed, as explained in Section 2.2.5.

**Table 2.** Summary of T2D1, T2D2, T2D3, T2DC datasets, and the numbers of identified SNPs, genes, subnetworks for each dataset.

| Datasets | # of Cases | # of Controls | # of SNPs | # of SNPs (p-value < 0.05) | # of rsIDs | # of Genes | # of Subnetworks |
| --- | --- | --- | --- | --- | --- | --- | --- |
| T2D1 | 12.931 | 57.196 | 14.683.492 | 762.111 | 335.212 | 15.806 | 984 |
| T2D2 | 62.892 | 596.424 | 5.053.015 | 557.564 | 557.564 | 15.460 | 904 |
| T2D3 | 74.124 | 824.006 | 21.635.866 | 1.525.650 | 639.622 | 17.800 | 941 |
| T2DC | - | - | - | - | - | 10.298 | 813 |

### 3.1 Affected subnetworks that are identified using meta GWAS datasets

For all datasets, the genes and their significance levels are mapped to protein-protein interaction network and 983, 903, 940 and 813 active protein subnetworks are identified for T2D1, T2D2, T2D3 and T2DC datasets, respectively. Numbers of the genes included in these subnetworks are depicted in Figure 3A for 70KforT2Dmeta-analysis dataset (T2D1), in Figure 3B for the meta-analysis of DIAGRAM, GERA, UKB GWAS datasets (T2D2), in Figure 3C for T2D3 dataset, in Figure 3D for T2DC dataset. While most of the subnetworks include 175-250 genes in T2D1 and T2D2 datasets, most of the subnetworks detected for T2C dataset include 200-250 genes. Around two third of the subnetworks, which are identified for T2D3 dataset include 150-175 genes. For each identified subnetwork, functional enrichment analysis is carried out and hence, affected pathways are determined.

**Figure 3.**
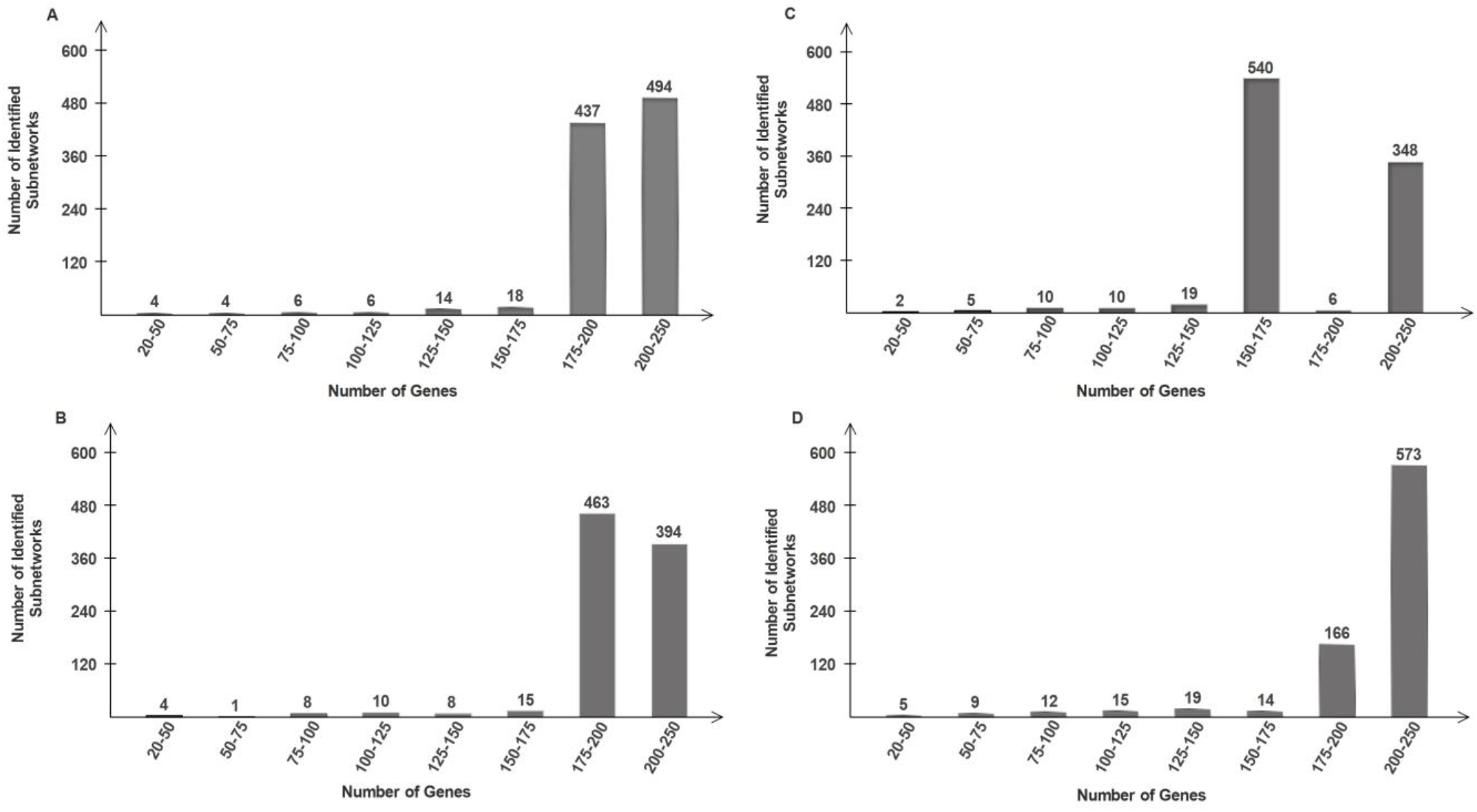
Numbers of genes included in the identified **(A)** 983 subnetworks for T2D1, **(B)** 903 subnetworks for T2D2, **(C)** 940 subnetworks for T2D3, and **(D)** 813 subnetworks for T2DC datasets.

### 3.2 Dysregulated modules of T2D that are identified using network propagation

Known T2D genes, collected in the (Ghiassian et al., 2015) study are used as seed genes to find dysregulated modules via expanding a module by adding other possible genes to the known disease gene clusters. This study indicated that seed proteins display unusual interaction patterns among each other. It enlightens the idea that the existence of disease specific modules is not by chance. Connectivity significance values are calculated for all neighbors of 73 known T2D disease associated seed genes. Afterwards, the node with the most significant interaction is added to the module and this iteration is repeated until 200 and 500 genes are included in a module. Then, functional enrichment procedure is performed on each of these two dysregulated modules (T2D_D200, T2D_D500).

### 3.3 Affected Pathways of T2D

Based on the observation that genes almost always act cooperatively rather than independently, to facilitate the biological interpretation of high-throughput data, many different methods have been postulated to identify the biological pathways associated with a particular clinical condition under study. Here, to characterize this cooperative nature of genes and to elucidate the molecular mechanisms of T2D, we investigate the affected pathways of T2D and search for the potential failures in these wiring diagrams.

#### 3.3.1 Overrepresented Pathways of T2D Dysregulated Modules

To detect possible pathogenic pathways related with T2D, the genes listed in each dysregulated module are compared with the genes included in KEGG pathways and the proportion of the module genes over all pathway-associated genes is calculated. Significantly affected KEGG pathways (pathways with corrected p-values < 0.05) for our defined dysregulated modules are appended to potentially significant pathway list of T2D disorder. Table 3 presents top 10 affected pathways that are found to be overrepresented in the dysregulated modules of T2DC dataset. Five of these pathways are also identified in all other T2D datasets. These shared pathways are Spliceosome, Focal adhesion, soluble N-ethylmaleimide-sensitive factor attachment protein receptor (SNARE) interactions in vesicular transport, transforming growth factor-β (TGF-β) signaling, and ErbB signaling pathways. Figures 4A and 4B depicts the commonalities among the top 50 and top 100 affected pathways enriched for the dysregulated modules of T2D1, T2D2, T2D3, T2DC datasets, and among the gold standard T2D pathways. As illustrated in Figure 4A, when the identified top 50 affected pathways are overlapped among all four datasets, 24 KEGG pathways are commonly observed. These pathways are Valine, leucine and isoleucine degradation, SNARE interactions in vesicular transport, Cholinergic synapse, TGF-beta signaling pathway, ErbB signaling pathway, Ubiquitin mediated proteolysis, Focal adhesion, ECM-receptor interaction, Gap junction, Spliceosome, Serotonergic synapse, Pathways in cancer, Retrograde endocannabinoid signaling, beta-Alanine metabolism, Neurotrophin signaling pathway, GABAergic synapse, Chemokine signaling pathway, Glioma, Dopaminergic synapse, Glutamatergic synapse, Endocytosis, GnRH signaling pathway, T cell receptor signaling pathway, Fc gamma R-mediated phagocytosis. When we compare these top 50 affected pathways of four datasets with the gold standard T2D pathway set (Yoon et al., 2018), Valine, leucine and isoleucine degradation pathway was commonly identified (as shown in Figure 4A). The comparison of the top 100 affected pathways of these datasets with gold standard T2D pathway set resulted in 8 common KEGG pathways, which are Valine, leucine and isoleucine degradation, Jak-STAT signaling pathway, Cell cycle, Glycolysis / Gluconeogenesis, Calcium signaling pathway, Insulin signaling pathway, Fatty acid metabolism, Wnt signaling pathway (as shown in Figure 4B).

**Figure 4.**
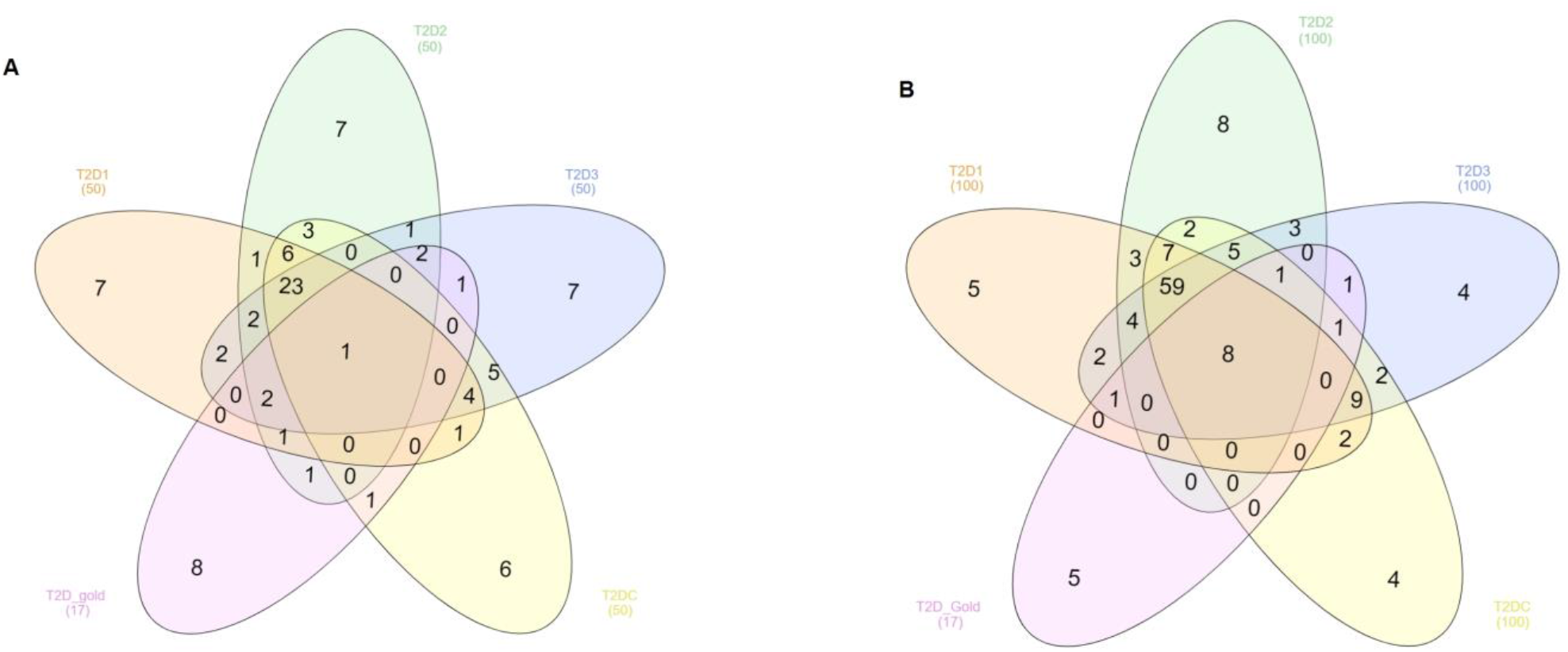
Commonalities between **(A)** top 50, and **(B)** top 100 affected pathways identified from T2D1, T2D2, T2D3, and T2DC datasets.

**Table 3.** Top 10 affected T2D pathways of T2DC dataset. Among these pathways, 5 pathways are commonly overrepresented for the dysregulated modules of T2D1, T2D2, T2D3, T2DC datasets.

|  | p-values |  |  |  | Rank |  |  |  | Number of genes |  | Percent of identified genes in associated pathways |
| --- | --- | --- | --- | --- | --- | --- | --- | --- | --- | --- | --- |
| KEGG term | T2DC | T2D1 | T2D2 | T2D3 | T2D C | T2D1 | T2D2 | T2D3 | T2D_Union | in pathways |  |
| Spliceosome | 8.55E-39 | 3.26E-27 | 6.95E-30 | 3.10E-41 | 1 | 15 | 8 | 5 | 104 | 127 | 0.81 |
| Focal adhesion | 7.032E-38 | 1.80E-30 | 3.82E-42 | 1.97E-54 | 2 | 10 | 1 | 1 | 172 | 200 | 0.86 |
| SNARE interactions in vesicular transport | 1.98E-35 | 1.37E-37 | 8.16E-33 | 5.41E-44 | 3 | 3 | 5 | 4 | 34 | 36 | 0.94 |
| Valine leucine and isoleucine degradation | 5.97E-35 | 3.26E-43 | 6.39E-20 | 3.34E-29 | 4 | 1 | 34 | 13 | 41 | 44 | 0.93 |
| Purine metabolism | 7.60E-34 | 5.35E-43 | 4.92E-12 | 1.29E-45 | 5 | 2 | 83 | 3 | 99 | 166 | 0.59 |
| Dopaminergic synapse | 3.26E-33 | 1.04E-20 | 9.48E-32 | 6.80E-34 | 6 | 37 | 7 | 9 | 119 | 130 | 0.91 |
| TGF-beta signaling pathway | 5.03E-29 | 8.70E-32 | 5.61E-34 | 3.23E-28 | 7 | 6 | 3 | 15 | 75 | 84 | 0.89 |
| ErbB signaling pathway | 1.59E-28 | 4.64E-31 | 1.00E-29 | 1.46E-37 | 8 | 8 | 9 | 7 | 85 | 87 | 0.97 |
| Chemokine signaling pathway | 5.23E-28 | 1.47E-21 | 1.01E-23 | 2.97E-19 | 9 | 33 | 20 | 39 | 163 | 189 | 0.86 |
| Glutamatergic synapse | 3.47E-27 | 1.97E-20 | 1.94E-29 | 3.03E-28 | 10 | 38 | 10 | 14 | 101 | 126 | 0.80 |

#### 3.3.2 Enriched Pathways for the Expanded Modules of T2D Seed Genes

Overrepresented pathways for expanded modules of 73 T2D seed genes, including 200 and 500 genes are identified with functional enrichment analysis. As shown in Table 4, the enrichment operation on T2D_D200 and T2D_D500 dysregulated modules resulted in 41 and 84 significant pathways, respectively.

**Table 4.**
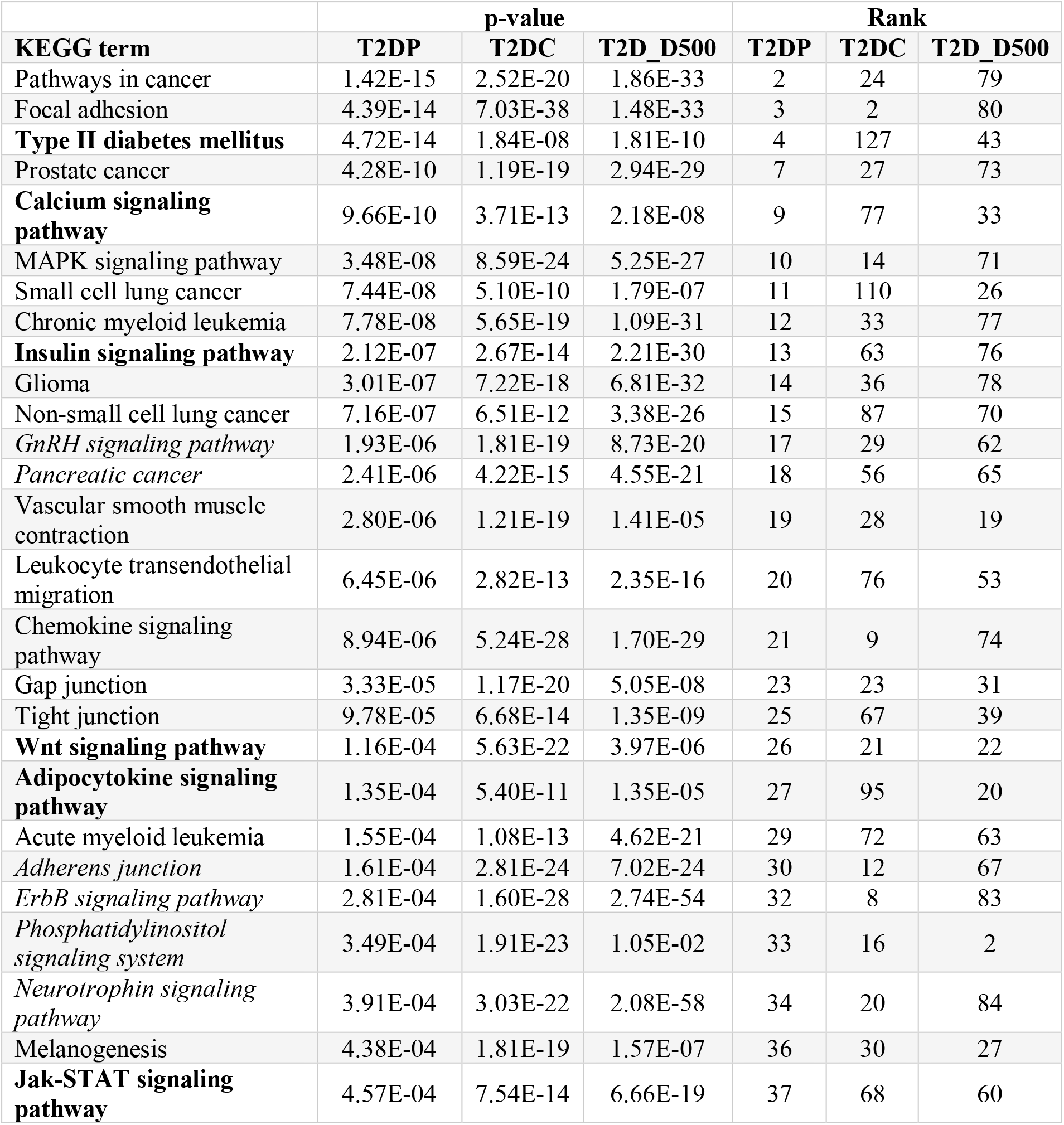
Comparison of the overrepresented pathways of T2D dysregulated modules (T2DC), expanded modules of T2D seed genes (T2D_D500), and the affected pathways identified using pathway scoring algorithm (T2DP).

#### 3.3.3 The Pathways that are Identified Using Pathway Scoring Algorithm on T2D GWAS meta data

The pathway scoring algorithm, as explained in Section 2.2.10 is used to find potentially affected pathways for T2D1, T2D2 and T2D3 data sets. Firstly, gene and pathway scores from SNP-phenotype association summary statistics are computed. Secondly, the calculated scores of affected pathways for each datasets are combined with Fisher’s method, and consequently, 38 KEGG and 46 Reactome pathways are detected for this combined data (T2D_PC).

In Table 4, the commonly identified KEGG pathways of T2DC, T2D_D500, T2D_PC methods, which are described in Sections 3.3.1, 3.3.2, 3.3.3, respectively, are listed. The affected pathways, which are highlighted in bold, refers to the gold standard KEGG pathways reported in the (Yoon et al., 2018)’s study. The affected pathways, which are highlighted in italic, refers to the pathways that are known in literature as related with T2D development mechanisms, as discussed in detail in Section 4. Among the 17 gold standard KEGG pathways of T2D, Type II diabetes mellitus, Calcium signaling, Insulin signaling, Wnt signaling, Adipocytokine signaling, and Jak-STAT signaling pathways are found with our methodology.

### 3.4 Shared T2D Subnetworks and Pathways Among Different GWAS meta data

#### 3.4.1 Comparative Evaluation of Identified T2D Subnetworks for Each Dataset

The identified T2D1, T2D2, T2D3 and T2DC subnetworks (as explained in Section 3.1, and summarized in Figure 3) are compared in a pairwise manner to assess the shared information among them. Firstly, for each x, y pairs of T2D1, T2D2, T2D3 and T2DC datasets, each identified subnetwork of T2D_x_ dataset and T2D_y_ dataset are compared in gene level and a contingency table of T2D_x_/T2D_y_, as shown in Table 1, is created. In this contingency table, each value of n_ij_ represents the shared gene counts between the i^th^ subnetwork of T2D_x_ dataset and the j^th^ subnetwork of T2D_y_ dataset. Secondly, based on this table, the entropy values H(T2D_x_), H(T2D_y_) and the mutual information values I(T2D_x_, T2D_y_) are computed for each x, y dataset pair. Thirdly, normalized MI is calculated as explained in Section 2.2.10. This procedure is repeated for all pairwise combinations of the T2D datasets. Hence, similarity scores (NMI_SUM_) are calculated between all pairs of datasets. The presented heatmap in Figure 5 illustrate the similarities of datasets according to the strength of the NMI_SUM_ score. As illustrated in Figure 5A, T2D1, T2D2, T2D3 and T2DC subnetwork similarities are resulted in range [0, 0.01]. While highest similarity score of 0.0073 is obtained for T2D2-T2D3 dataset pair, the lowest score of 0.0060 is obtained for T2D1-T2DC dataset pair. Accordingly, while the darker colors indicate higher correlation, lighter colors indicate smaller correlation in the heatmap of Figure 5. In Figure 5, NMI_SUM_ scores in the diagonals of the heatmap are “whitened” for clearer visibility of the other NMI_SUM_ values.

**Figure 5.**
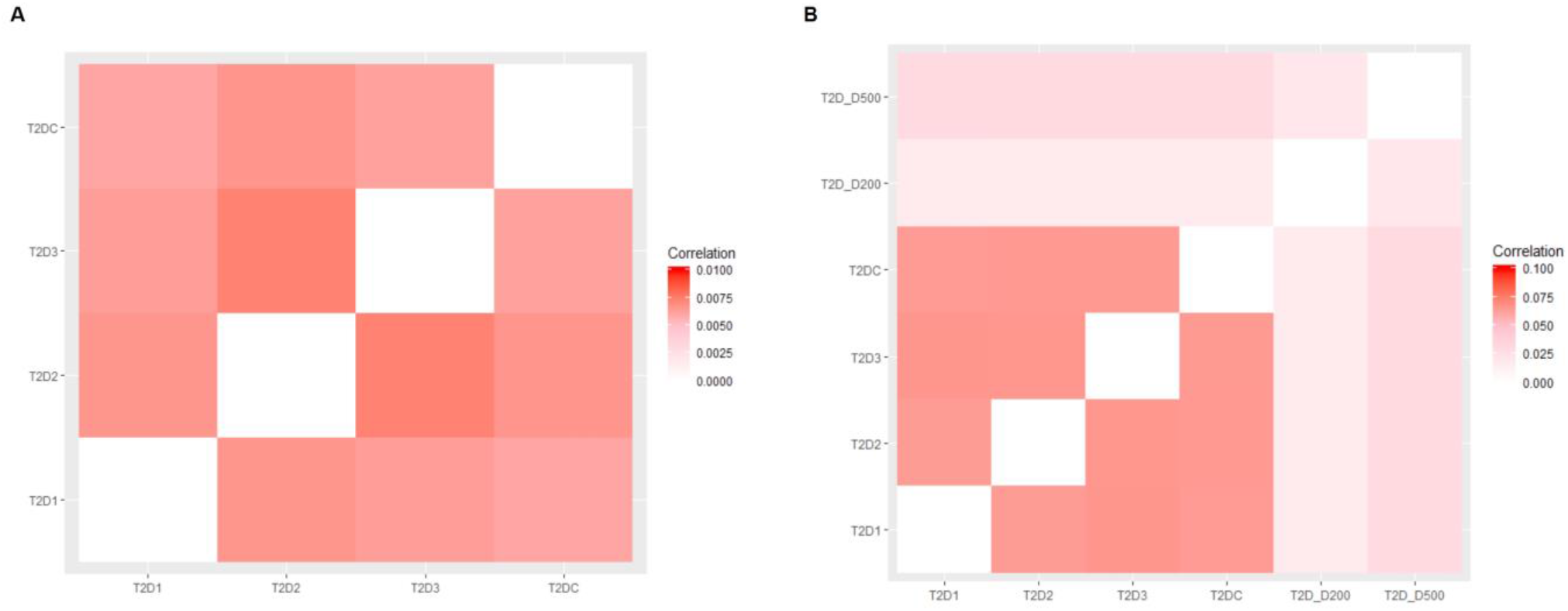
Shared information comparison among different datasets in terms of **(A)** identified T2D subnetworks, and **(B)** identified pathways via normalized mutual information (NMI_SUM_). While the darker colors indicate higher correlation, lighter colors indicate smaller correlation. NMI_SUM_ scores in the diagonals of the heatmap are “whitened” for clearer visibility of the other NMI_SUM_ values.

#### 3.4.2 Comparative Evaluation of Identified T2D Pathways for Each Dataset

Shared information among different methodologies (subnetwork identification, as presented in Section 2.2.5.1 and bottom-up approach, as presented in Section 2.2.5.2) and different T2D meta-datasets, are also evaluated in terms of the identified T2D pathways. The same functional enrichment analysis is applied on the subnetworks and dysregulated modules, as explained in Section 2.2.6. In addition to the identified pathways of T2D1, T2D2, T2D3 and T2DC datasets, the pathways identified from T2D_D200 and T2D_D500 gene sets are also evaluated here. Firstly, for each x, y pairs of T2D1, T2D2, T2D3, T2DC, T2D_D200 and T2D_D500, each identified pathway of T2D_x_ dataset and T2D_y_ dataset are compared in terms of their common genes and a contingency table of T2D_x_/T2D_y_ is created, as shown in Table 1. In this contingency table, each value of n_ij_ represents the shared gene counts between the i^th^ identified pathway of T2D_x_ dataset and the j^th^ identified pathway of T2D_y_ dataset. Secondly, based on this table, the entropy values H(T2D_x_), H(T2D_y_) and mutual information values I(T2D_x_, T2D_y_) are computed for each x, y dataset pair. Thirdly, normalized MI is calculated as explained in Section 2.2.10. This procedure is repeated for all pairwise combinations of the T2D datasets. Hence, similarity scores (NMI_SUM_) are calculated between all pairs of datasets, in terms of overrepresented pathways. In terms of the identified pathways, Figure 5B illustrates the similarity levels of the T2D1, T2D2, T2D3, T2DC, T2D_D200 and T2D_D500, in the range of [0-0.1]. While a maximum NMI_SUM_ score of 0.0658 is achieved for T2D1-T2D3 pair, a minimum NMI_SUM_ score of 0.016 is obtained for T2DC-T2D_D200 pair.

### 3.5 Affected Pathway Subnetworks and Pathway Clusters of T2D

We hypothesized that similar to the dysregulated modules of proteins, dysregulated modules of pathways have a role in disease development mechanisms. In order to identify affected pathway subnetworks of a disease; we proposed a methodology, as shown in Figure 2. Instead of a PPI network, this method requires a pathway network as the baseline. Here, we utilized the 288 human KEGG pathways as a reference, for the generation of this biological network. To establish a pathway network, firstly, we examined the relationships between the genes and the biological pathways, as explained in Section 2.2.7. In this study, we stored these relationships in a gene-term matrix, which is a binary matrix with dimensions 6881 * 288, representing the number of individual genes in all pathways, and the number of pathways, respectively. Secondly, the relationships between the pathways are analyzed, as explained in Section 2.2.7. For this purpose, kappa statistics was used to determine the relationships between pathways, and a term-term matrix (of size 288 *288), was formed by using the previously obtained gene-term matrix. Thirdly, to identify interrelated pathways, we experimented with different cutoff values of kappa scores. The sizes of the networks that are created with different threshold values are presented in Table 5. Since the node to edge ratio in the human PPI network is approximately 1 to 10, the kappa score threshold value is selected as 0.15 in this study and finally, a human pathway network including 288 pathways (nodes) and 2976 interrelations (edges) is created.

**Table 5.** Node – Edge relationships in the generated pathway networks and affected pathway subnetworks.

| Sizes of the generated pathway networks for different threshold values |  |  |
| --- | --- | --- |
| Threshold Values ( ≥ ) | # of Nodes | # of Edges |
| 0 | 288 | 82944 |
| 1.21E-5 | 288 | 10904 |
| 0.05 | 288 | 6806 |
| 0.1 | 288 | 4617 |
| 0.15 | 288 | 2976 |
| 0.2 | 288 | 1866 |
| 0.25 | 288 | 1321 |

**Sizes of the generated highest scoring pathway subnetworks for different T2D datasets**
| Dataset | # of Nodes | # of Edges |
| --- | --- | --- |
| T2D1 | 119 | 1356 |
| T2D2 | 134 | 1383 |
| T2D3 | 135 | 1441 |
| T2DC | 158 | 1709 |

Active subnetwork identification algorithms require a biological network and the significance values of the nodes, e.g. the p-values of the genes obtained from microarray studies, indicating the significance of a gene, in terms of the expression levels differing between two experimental conditions. Here, while our biological network is selected as our generated pathway network, significance values of the nodes are selected as the corrected hypergeometric test p-values, indicating the importance of the pathway for T2D. Following the methodology proposed in Figure 2, for all T2D datasets, only one affected pathway subnetwork exceeded the predefined subnetwork score, as summarized in Table 5. As the node and edge numbers of these identified pathway subnetworks could be inspected from Table 5, it could be observed that the nodes are severely connected to each other in the identified pathway subnetworks. Therefore, these four identified pathway subnetworks (for four different datasets) were further grouped into subcategories as explained in Section 2.2.8, and the affected pathway clusters of T2D are obtained for each dataset. As shown in Table 6, for T2D1, T2D2, T2D3, T2DC datasets, 7, 9, 7, and 8 affected pathway clusters are identified respectively. Numbers of nodes (pathways) included in each cluster and the scores of each pathway cluster can be found in Table 6. When the obtained results are analyzed, it is seen that the initial pathway subnetwork, which is severely connected with each other and has more than 50 nodes is successfully divided into smaller disease related subnetworks. This can be considered as a proof of the effectiveness of the developed method. The highest scoring pathway cluster of T2D1, T2D2, T2D3, T2DC datasets included 38, 34, 35 and 35 pathways, respectively. For each dataset, the representative networks of the identified pathway clusters are shown in Figure 6. When we analyze the commonalities among these pathways, we observed in Figure 7 that 29 of these pathways were commonly identified in T2D1, T2D2, T2D3, T2DC datasets. The details of these commonly identified pathways are given in Table 7.

**Figure 6.**
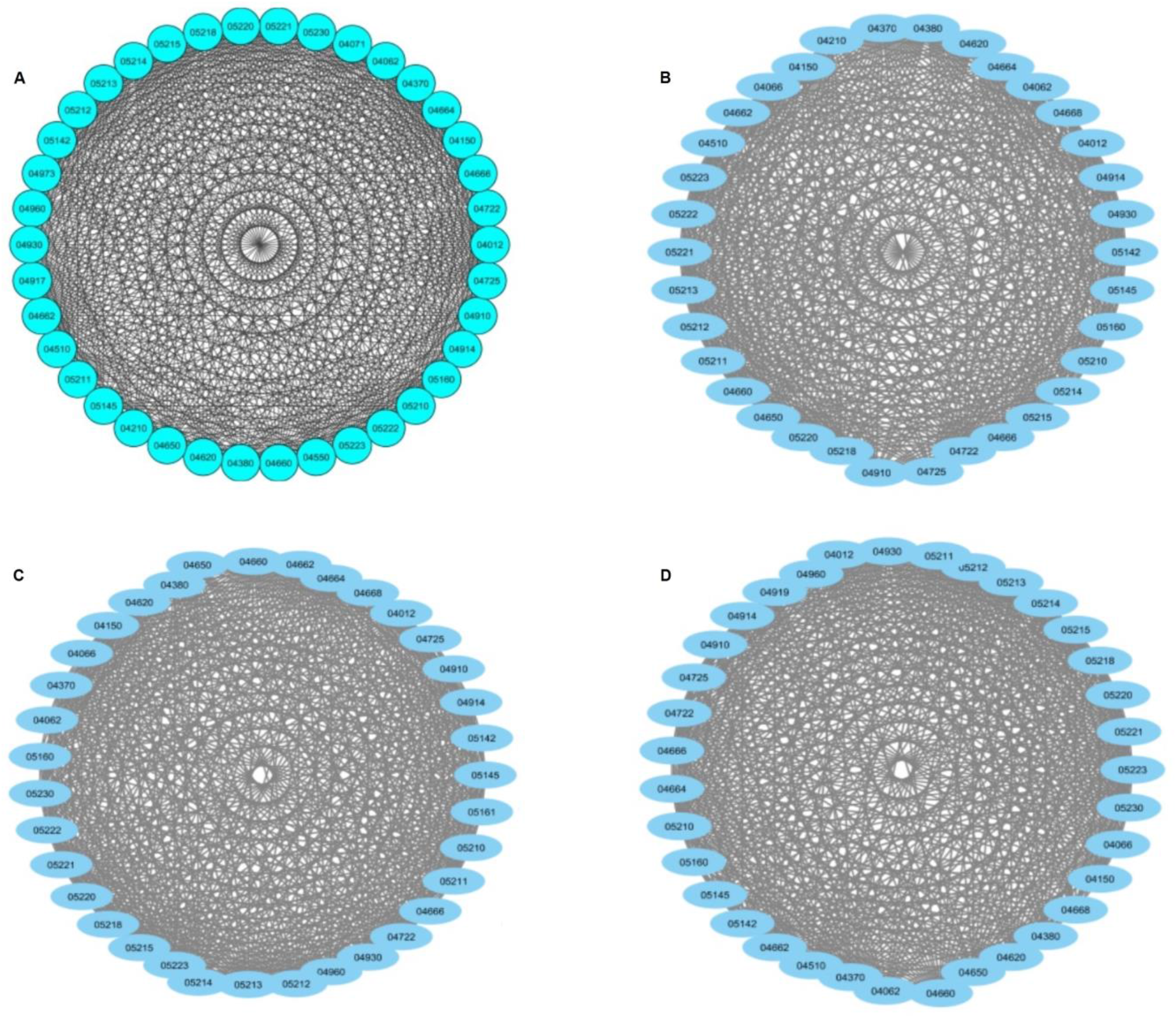
The representative networks of the highest scoring pathway clusters of **(A)** T2D1, **(B)** T2D2, **(C)** T2D3, **(D)** T2DC datasets, including 38, 34, 35 and 35 pathways, respectively.

**Figure 7.**
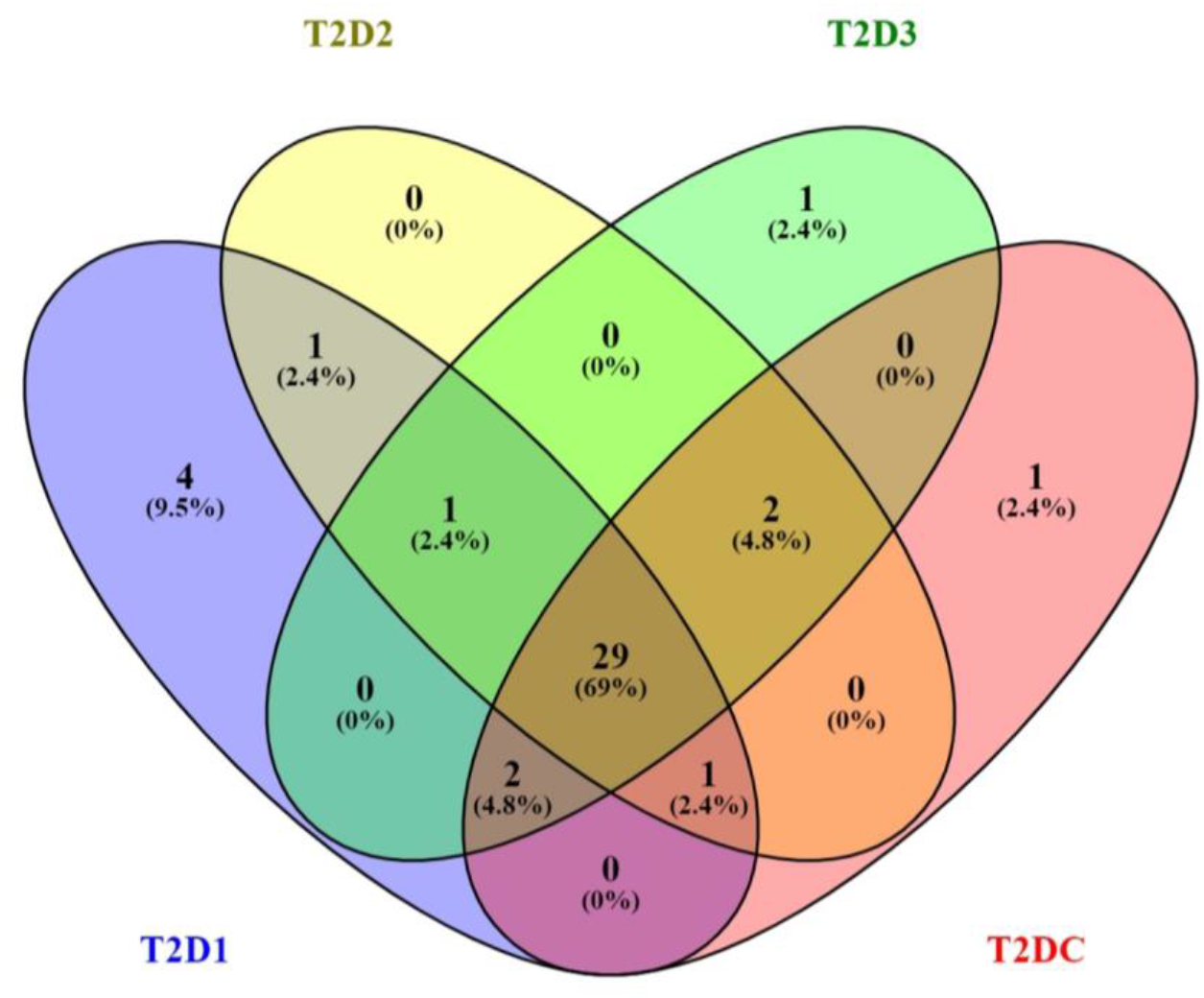
Commonalities between the highest scoring pathway clusters of T2D1, T2D2, T2D3, and T2DC datasets.

**Table 6.** Identified pathway clusters that are affected in T2D for each dataset.

| T2D1 |  |  | T2D2 |  |  | T2D3 |  |  | T2DC |  |  |
| --- | --- | --- | --- | --- | --- | --- | --- | --- | --- | --- | --- |
| # of Clusters | # of Nodes | Score of Cluster | # of Clusters | # of Nodes | Score of Cluster | # of Clusters | # of Nodes | Score of Cluster | # of Clusters | # of Nodes | Score Of Cluster |
| 7 | 38 | 32.919 | 9 | 34 | 30.182 | 7 | 35 | 31.412 | 8 | 35 | 31.118 |
|  | 14 | 8.462 |  | 19 | 13.111 |  | 21 | 14.3 |  | 16 | 8.8 |
|  | 9 | 4.75 |  | 15 | 5.286 |  | 11 | 5.2 |  | 16 | 8.533 |
|  | 4 | 3.333 |  | 5 | 5 |  | 5 | 4.5 |  | 11 | 5 |
|  | 3 | 3 |  | 5 | 4,5 |  | 4 | 4 |  | 5 | 5 |
|  | 3 | 3 |  | 4 | 4 |  | 4 | 3.333 |  | 8 | 4.286 |
|  | 3 | 3 |  | 3 | 3 |  | 3 | 3 |  | 4 | 3.333 |

\*Cut Off Value: 0.2, Haircut: True Fluff: FALSE, K-Core: 2

**Table 7.** Common pathways of highest scoring pathway clusters that are identified for different T2D GWAS meta-data.

| Pathway Name | p-values |  |  |  | Rank |  |  |  |
| --- | --- | --- | --- | --- | --- | --- | --- | --- |
|  | T2D1 | T2D2 | T2D3 | T2DC | T2D1 | T2D2 | T2D3 | T2DC |
| Renal cell carcinoma | 7.12E-15 | 1.95E-15 | 7.23E-13 | 8.14E-15 | 68 | 55 | 90 | 57 |
| Colorectal cancer | 1.52E-12 | 7.53E-10 | 1.82E-14 | 3.51E-17 | 97 | 115 | 77 | 41 |
| Hepatitis C | 2.99E-14 | 1.29E-14 | 1.35E-18 | 1.59E-16 | 77 | 62 | 47 | 43 |
| VEGF signaling pathway | 1.05E-11 | 1.20E-10 | 4.18E-12 | 4.15E-13 | 104 | 99 | 99 | 78 |
| Toxoplasmosis | 2.38E-12 | 2.24E-12 | 1.30E-18 | 4.39E-13 | 99 | 78 | 48 | 80 |
| Chagas disease (American trypanosomiasis) | 2.10E-18 | 1.62E-12 | 3.85E-19 | 3.57E-15 | 48 | 76 | 42 | 54 |
| Type II diabetes mellitus | 1.32E-12 | 2.68E-09 | 6.18E-19 | 1.84E-08 | 96 | 124 | 44 | 127 |
| Chemokine signaling pathway | 1.47E-21 | 1.01E-23 | 2.97E-19 | 5.23E-28 | 33 | 20 | 39 | 9 |
| Progesterone-mediated oocyte maturation | 2.67E-16 | 3.57E-12 | 4.95E-16 | 7.25E-18 | 62 | 81 | 68 | 37 |
| Insulin signaling pathway | 2.16E-16 | 1.67E-16 | 2.96E-18 | 2.67E-14 | 60 | 48 | 49 | 63 |
| Toll-like receptor signaling pathway | 1.70E-29 | 2.63E-11 | 3.20E-13 | 1.27E-14 | 13 | 91 | 85 | 62 |
| Cholinergic synapse | 6.32E-35 | 1.17E-25 | 1.61E-31 | 4.37E-27 | 4 | 16 | 11 | 11 |
| Neurotrophin signaling pathway | 4.20E-22 | 3.68E-23 | 3.03E-31 | 3.02E-22 | 30 | 22 | 12 | 20 |
| Fc gamma R-mediated phagocytosis | 3.57E-19 | 2.88E-18 | 1.01E-19 | 1.75E-16 | 44 | 37 | 35 | 47 |
| Osteoclast differentiation | 5.24E-22 | 1.28E-14 | 3.60E-19 | 3.16E-17 | 31 | 61 | 41 | 40 |
| T cell receptor signaling pathway | 3.32E-19 | 3.69E-21 | 4.49E-20 | 2.14E-18 | 43 | 32 | 33 | 34 |
| Fc epsilon RI signaling pathway | 3.75E-18 | 9.42E-16 | 5.92E-18 | 2.33E-23 | 52 | 53 | 52 | 17 |
| Natural killer cell mediated cytotoxicit | 2.61E-13 | 1.53E-13 | 2.12E-09 | 5.47E-12 | 90 | 69 | 131 | 86 |
| B cell receptor signaling pathway | 3.28E-19 | 3.39E-17 | 2.41E-14 | 1.96E-19 | 42 | 43 | 78 | 31 |
| mTOR signaling pathway | 1.28E-12 | 4.34E-10 | 1.72E-08 | 1.60E-10 | 95 | 108 | 141 | 102 |
| Non-small cell lung cancer | 7.60E-16 | 3.04E-11 | 1.86E-13 | 6.51E-12 | 65 | 92 | 82 | 87 |
| ErbB signaling pathway | 4.64E-31 | 1.09E-29 | 1.46E-37 | 1.59E-28 | 8 | 9 | 7 | 8 |
| Acute myeloid leukemia | 5.42E-14 | 1.40E-10 | 1.03E-11 | 1.08E-13 | 80 | 102 | 105 | 72 |
| Chronic myeloid leukemia | 7.27E-20 | 8.58E-17 | 2.48E-16 | 5.65E-19 | 41 | 45 | 65 | 33 |
| Melanoma | 4.79E-14 | 8.51E-17 | 6.46E-15 | 1.05E-14 | 78 | 44 | 74 | 59 |
| Prostate cancer | 1.13E-17 | 1.82E-13 | 1.12E-12 | 1.18E-19 | 53 | 70 | 93 | 27 |
| Glioma | 3.33E-21 | 1.67E-16 | 7.34E-19 | 7.21E18 | 35 | 47 | 45 | 36 |
| Endometrial cancer | 3.47E-16 | 1.67E-14 | 4.80E-13 | 1.62E16 | 63 | 63 | 88 | 45 |
| Pancreatic cancer | 6.15E-13 | 4.15E-14 | 8.21E-15 | 4.21E-15 | 94 | 65 | 75 | 56 |

## 4 Discussion

GWASs of T2D have significantly accelerated the discovery of T2D–associated loci (Adeyemo et al., 2015; Bonnefond and Froguel, 2015; Liu et al., 2017; Meyre, 2017; Scott et al., 2017). Although the identified T2D-risk variants including 243 loci and 403 distinct association signals exhibit a potential for clinical translation, the genome-wide chip heritability explains only 18% of T2D risk (Bonàs-Guarch et al., 2018; Mahajan et al., 2018a; Xue et al., 2018). Traditional GWASs focus on top-ranked SNPs and discard all others except ‘the tip of the iceberg’ SNPs. Such GWAS approaches are only capable of revealing a small number of associated functions. In this regard, even though GWASs are a compelling method to detect disease-associated variants, it does not directly address the biological mechanisms underlying genetic association signals, and hence, the development of novel post-GWAS analysis methodologies is needed (Lin et al., 2017), (Gallagher and Chen-Plotkin, 2018), (Erdmann and Zeller, 2019). In this respect, to enlighten the molecular mechanisms of Type 2 Diabetes development, here we proposed a method that perform protein subnetwork, pathway subnetwork and pathway cluster level analyses of the SNPs that are found to be mildly associated with T2D in multiple association studies. In other words, to achieve a coherent comprehension of T2D molecular mechanisms, the proposed network and pathway-based solution conjointly analyzes three meta-analyses of GWAS, which are conducted on T2D.

The baseline of our study is built on the interactions of T2D related proteins, since the proteins act as the functional base units of the cells and construct the frameworks of cellular mechanisms. Protein network structure helps us to gain a collective insight about the biological systems. At the mesoscopic level of these protein networks, active modules are the potential intermediate building blocks between individual proteins and the global interaction network. Dysregulation of these modules are considered to have a role in disease development mechanisms. Hence, the identification of dysregulated modules of T2D helps us to understand the fundamental molecular characteristics of T2D and to discover new candidate disease genes having a role in the regulation of T2D related pathways. In this context, for each analyzed T2D GWAS meta-analysis dataset, where the characteristics of each dataset is summarized in Table 2, 800 to 1000 dysregulated modules, including 150 to 250 genes are detected using a top-down approach, as explained in Section 2.2.5.1. As outlined in Figure 1, these modules are functionally enriched and the pathways that have a potential effect on T2D development are identified. As presented in Table 3, among the top 10 affected T2D pathways of T2DC datasets, 5 pathways are commonly overrepresented for the dysregulated modules of T2D1, T2D2, T2D3, T2DC datasets. These five shared pathways are Spliceosome, Focal adhesion, SNARE interactions in vesicular transport, TGF-β signaling, and ErbB signaling pathways. All these pathways are known to have a role in T2D development mechanisms. Spliceosome pathway has a role in the regulation of alternative splicing in insulin resistance cases by aberrantly spliced genes like ANO1, GCK, SUR1, VEGF (Costantini et al., 2011; Dlamini et al., 2017; Schmid et al., 2012). Focal adhesion pathway is complementary in regulation of insulin signaling pathway. Via controlling adipocyte survival, Focal adhesion kinases (FAK) regulate insulin sensitivity (Luk et al., 2017). SNARE protein contributes to fusion mechanism of insulin secretory vesicles (Xiong et al., 2017). The study conducted by Boström *et. al.* demonstrated that total skeletal muscle SNARE protein SNAP23 and SNARE related Munc18C protein levels are higher in patients with type 2 diabetes, which are also correlated with markers of insulin resistance (Boström et al., 2010). TGF-β signaling pathway has role in inflammation by cytokines such as interleukins, tumor necrosis factors, chemokins interferons, transforming growth factors (TGF). Insulin enhances TGF-β receptors in fibroblasts and epithelial cells. Herder *et. al.* documented that high levels of anti-inflammatory immune mediator TGF-β1 are correlated with T2D (Herder et al., 2009). TGF-β signaling pathway is also shown to have a crucial role in extracellular matrix accumulation in diabetic nephropathy (Kajdaniuk et al., 2013). Akhtar *et. al.* showed that the dysregulation of epidermal growth factor receptor family (ErbB) triggers vascular dysfunction stimulated by hyperglycemia in T2D (Akhtar et al., 2015). Other dual role of ErbB protein family included diabetes triggered cardiac dysfunction (Akhtar and Benter, 2013).

While identifying active subnetworks of T2D, in addition to the top-down approach (as discussed above), we also applied bottom-up approach as explained in Section 2.2.5.2. Overrated pathways of i) top-down approach (T2DC), ii) bottom-up approach (T2D_D200, T2D_D500), and iii) pathway scoring algorithm (T2D_P) are comparatively evaluated. Among these pathways, Type II diabetes mellitus, Calcium, Insulin, Wnt, Adipocytokine, Jak-STAT signaling pathways (shown in bold in Table 4) overlap with gold standard pathways of T2D (Yoon et al., 2018). Additionally, the pathways that are shown in italic in Table 4, have support from the literature as following. The study conducted by (Berntorp et al., 2013) reported that T2D patients express antibodies against gonadotropin-releasing hormone GnRH in serum. (De Souza et al., 2016) stated T2D as prognostic and risk factor for pancreatic cancer. (Houtz et al., 2016) reported that paracrine neurotrophin signaling have a role in insulin secretion between pancreatic vascular system and beta cells, which is triggered by glucose. (Ono et al., 2001) stated that phosphatidylinositol signaling system including PTEN (phosphatase and tensin homologue deleted on chromosome 10) and PI3K (phosphoinositide3-kinase) proteins regulate glucose homeostasis and insulin metabolism. In a study performed by (Dissanayake et al., 2018), cadherin mediated adherens junction proteins are shown to have a potential regulation role in insulin secretion mechanism by controlling vesicle traffic in cell. Via studying different GWAS meta-analyses, Schierding *et. al.*, indicated the spatial connection of CELSR2–PSRC1 locus with BCAR3, which is part of the insulin signaling pathway (Schierding and O’Sullivan, 2015). The post-GWAS study conducted by (Liu et al., 2017) identified T2D risk pathways. Among these pathways, Type II diabetes mellitus, Calcium signaling pathway, Pancreatic cancer, MAPK signaling pathway, Chemokine signaling pathway, Tight junction pathways were also identified in our study (p<0.05). Another study performed by (Perry et al., 2009) analyzed T2D GWAS data and reported that Wnt signaling pathway, Olfactory transduction, Galactose metabolism, Pyruvate metabolism, Type II diabetes, TGF-signaling pathways are associated with T2D. Wnt signaling and Type II diabetes pathways are overlapped with our findings, as shown in Table 4. The analysis of T2D WTCCC GWAS dataset by (Zhong et al., 2010) indicated 22 affected pathways in T2D. Among these pathways, Tight junction, Phosphatidylinositol signaling system, Pancreatic cancer, Adherens junction, Calcium signaling pathway are replicated in our study, as shown in Table 4.

Using the mutual information based on the shared genes, the identified protein subnetworks and the affected pathways of each dataset were compared. While the NMI_SUM_ subnetwork scores range from 0 to 0.01, NMI_SUM_ pathway scores range from 0 to 0.1 (as shown in Figure 5). Hence, we show that while the subnetwork level analyzes increase the degree of irregularity, pathway level evaluation of different T2D GWAS meta-data and different methodologies (top-down vs. bottom-up approach) resulted in higher levels of conservation and yielded in more interpretable outcome.

While the Type II diabetes mellitus pathway is identified in the later rankings for T2D1, T2D2, T2D3, and T2DC GWAS datasets (as shown in Table 7), the incorporation of the generated pathway network information helped us to prioritize this pathway. This pathway is found in the highest scoring pathway cluster of each dataset. Since the pathways are strongly interrelated, our proposed approach created a pathway network, and identified affected pathway subnetworks and pathway clusters using multiple association studies, which are conducted on T2D. Our approach is based on both significance level of an affected pathway and its topological relationship with its neighbor pathways.

In conclusion, the availability of T2D GWAS meta-data and new analytical methods has provided opportunities to bridge the knowledge gap from sequence to consequence. In this study, the collective effects of T2D–associated variants are inspected using network and pathway-based approaches, and the prominent genetic association signals related with T2D biological mechanisms are revealed. We presented a comprehensive analysis of three different T2D GWAS meta-data at protein subnetwork, pathway, and pathway subnetwork levels. To explore whether our results recapitulate the pathophysiology of T2D, we performed functional enrichment analysis on the dysregulated modules of T2D. In addition to our analysis of the shared information among different datasets in terms of subnetworks, we also analyzed the shared information in terms of the identified T2D pathways. The identified pathway subnetworks, pathway clusters and affected genes within these pathways helped us to illuminate T2D development mechanisms. We hope the affected genes and variants within these identified pathway clusters help geneticists to generate mechanistic hypotheses, which can be targeted for large-scale empirical validation through massively parallel reporter assays at the variant level; and through CRISPR screens in appropriate cellular models, and through manipulation in in-vivo models, at the gene level.

## 6 Author Contributions

BBG and MUY conceived the ideas and designed the study. BBG, MUY, GG, MT conducted the experiments and analyzed the results. BBG, MUY, GG, and MT participated in the discussion of the results and writing of the article. All authors read and approved the final version of the manuscript.

## 7 Acknowledgments

We would like to thank the anonymous reviewers for their valuable comments and suggestions to improve the quality of the paper. We are also very grateful to Prof. David Torrents from Barcelona Supercomputing Center, to help us with the 70KforT2D meta-analysis data. We also would like to thank Prof. Albert-Laszlo ́ Barabasi at University of Notre Dame and Dr. Michael Cusick at Center for Cancer Systems Biology for providing us PPI dataset; Dr. Gabriela Bindea from Integrative Cancer Immunology Team of Cordeliers Research Center for her help with the ClueGO tool.

